# Lack of Fzd6 in Ciliated Cells Suppresses Ferroptotic Pulmonary Alveolar Cell Death Induced by LPS and Coronavirus

**DOI:** 10.1101/2023.01.23.524390

**Authors:** Qianying Yuan, Yi Luan, Barani Kumar Rajendran, Susan Compton, Wenwen Tang, Dianqing Wu

## Abstract

Pulmonary inflammation compromises lung barrier function and underlies many lung diseases including acute lung injury and acute respiratory distress syndrome (ARDS). However, mechanisms by which lung cells respond to the damage caused by the inflammatory insults are not completely understood. Here we show that Fzd6-deficiency in Foxj1^+^ ciliated cells reduces pulmonary permeability, lipid peroxidation, and alveolar cell death accompanied with an increase in alveolar number in lungs insulted by LPS or a mouse coronavirus. Single-cell RNA sequencing of lung cells indicates that the lack of Fzd6, which is expressed in Foxj1^+^ cells, increases expression of the aldo-keto reductase Akr1b8 in Foxj1^+^ cells. Intratracheal administration of the Akr1b8 protein phenocopies Fzd6-deficient lung phenotypes. In addition, ferroptosis inhibitors also phenocopy Fzd6-deficient lung phenotypes and exert no further effects in Fzd6-deficient lungs. These results reveal an important mechanism for protection of alveolar cells from ferroptotic death during pulmonary inflammation by Foxj1^+^ ciliated cells via paracrine action of Akr1b8.

## INTRODUCTION

Bacterial and viral infection of lungs causes acute pulmonary inflammation, which can lead to acute lung injury (ALI) and its more severe form acute respiratory distress syndrome (ARDS). ALI/ARDS are characterized by diffuse alveolar damage, neutrophilic inflammation, and increased permeability of the alveolar-capillary barrier, leading to protein-rich edema and subsequent impairment of arterial oxygenation [1-3]. ALI/ARDS represents a serious health problem with a high mortality rate [4, 5]. The incidence of ALI/ARDS is reported to be around 200,000 per year in the US with a mortality rate of around 40% before the current COVID-19 pandemic [4]. The recent COVID-19 as well as previous SARS-CoV and Middle Eastern respiratory syndrome (MERS) are all associated with respiratory illness due to alveolar damage, which can lead to severe ALI/ARDS and progressive respiratory failure and contribute to high mortality rates of these diseases [6-8]. Currently there are no pharmacological interventions for ALI/ARDS. The incomplete understanding of ALI/ARDS pathogenesis and pulmonary cell responses to inflammation hampers the progress in ALI/ARDS therapeutic development [9, 10].

Ferroptosis is a form of cell death caused by iron-dependent lipid peroxidation. Mechanistically, oxygen and iron, which are essential drivers of metabolism, cause the production of reactive species (ROS) and subsequent production of lipid peroxides. If these lipid peroxides cannot be neutralized efficiently and subsequently accumulate, they disrupt cellular membrane integrity and result in cell death. Ferroptosis plays important roles in many diseases including cancer, neurodegeneration, and ischemic organ injury [11-14]. It is also implicated in lung diseases as lipid peroxidation and effects of ferroptotic inhibitors were detected [13, 15-20]. However, it is unclear if ferroptotic cells actually occur during acute lung inflammation.

Wnt signaling regulates a wide range of physiological and pathophysiological processes. The best characterized Wnt signaling pathway is the Wnt-β-catenin pathway, in which Wnt proteins bind to their cell surface receptor Frizzled (Fzd) and LDL-related protein 5/6 (LRP5/6), leading to β-catenin stabilization and β-catenin-mediated transcription. There are 10 Fzd genes in mammals. Although many of them have been shown to play roles in various developmental processes, their roles in physiological and pathophysiological processes remain understudied. In this study, we found that the loss of Fzd6 in ciliated cells enhanced pulmonary barrier function and protected ferroptotic death of alveolar cells during lung inflammation induced by LPS or MHV, a mouse coronavirus that produces pulmonary pathological features of SARS [21]. This study revealed a previously unknown interaction of ciliated cells with alveolar cells in acute lung inflammation.

## RESULTS

### Fzd6 KO reduces lung permeability and increases alveolar number

Examination of Fzd6 tissue expression using the FANTOM 5 database [22] revealed that the lung was one of the top-ranked among the human tissues where Fzd6 is highly expressed (Fig. S1A). We investigated whether Fzd6 KO had a role in LPS-induced lung injury using a MX1-Cre-driven inducible Fzd6 KO mouse line and found that Fzd6 KO reduced protein contents in the bronchoalveolar lavage (BAL) and pulmonary permeability of mice received intratracheal administration of LPS for 18 hours compared to the wildtype littermate controls (Figure 1A). Pulmonary permeability was determined by measuring the FITC-BSA content in BAL from mice receiving I.V. infusion of FITC-BSA two hours before BAL collection. There were no differences in the BAL concentrations of IL-6, TNFα, and IL-1β (Fig. S1B) or in myeloid cell abundance in the BAL and lungs between WT and Fzd6 KO mice (Fig. S1C-D). Analysis of lung histomorphometry showed no significant difference in septal thickness, but revealed an increase in the number of alveoli in the Fzd6 KO lungs over the controls (Fig. 1B-D).

**Figure 1.**
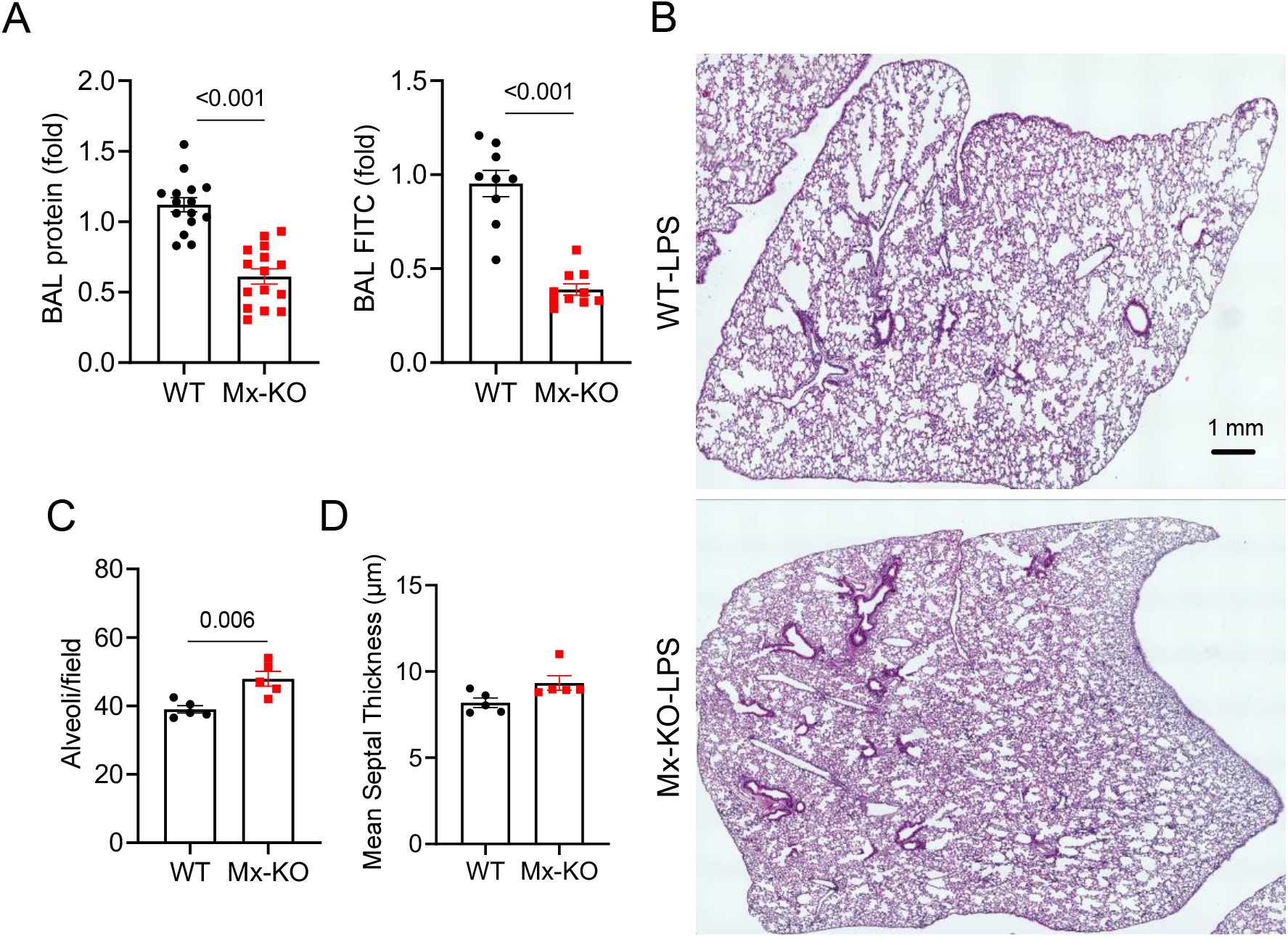
Fzd6 KO alters responses to pulmonary inflammation induced by LPS. Mx-Cre Fzd6 KO (Mx-KO) mice and their wildtype (WT) controls were treated with poly IC. They were then given LPS intratracheally one month later. The mice were administrated with FITC-BSA 18 hours after LPS treatment. Two hours later, the mice were euthanized, and protein and FITC-BSA contents in the BALs were measured (**A**). Some of the lung tissues were sectioned and stained with H&E, followed with quantification (**B**-**D**). Representative lung section scan images are shown in **B**. Each datum point represents one mouse. Data are presented as means±sem (two-tailed Student’s t-test).

### Fzd6 KO in Foxj1^+^ cells reduces lung permeability and increases alveolar number

The analysis of single-cell RNA sequencing data of CD45^+^ and CD45^-^ cells sorted from LPS-injured lungs revealed that Fzd6 was primarily expressed in CD45^-^ cells, more specifically Cdh5^+^ endothelial and Epcam^+^ epithelial cells, but not in CD45^+^ cells (Fig. 2A-B). Among the epithelial cells, the Foxj1^+^ ciliated cells expressed Fzd6 at a higher level than other CD45^-^ cells (Fig. S2A). To determine the cell type in which Fzd6 KO contributes to the resistance to LPS-induced lung injury, we generated endothelial cell-specific (driven by Cdh5-Cre) and Foxj1^+^ cell-specific (driven by Foxj1-Cre) Fzd6 KO mouse lines, respectively. The mice lacking Fzd6 in Foxj1^+^ cells (Fig. 2C), but not those lacking Fzd6 in endothelial cells (Fig. 2D), showed increased resistance to LPS-induced lung permeability.

**Figure 2.**
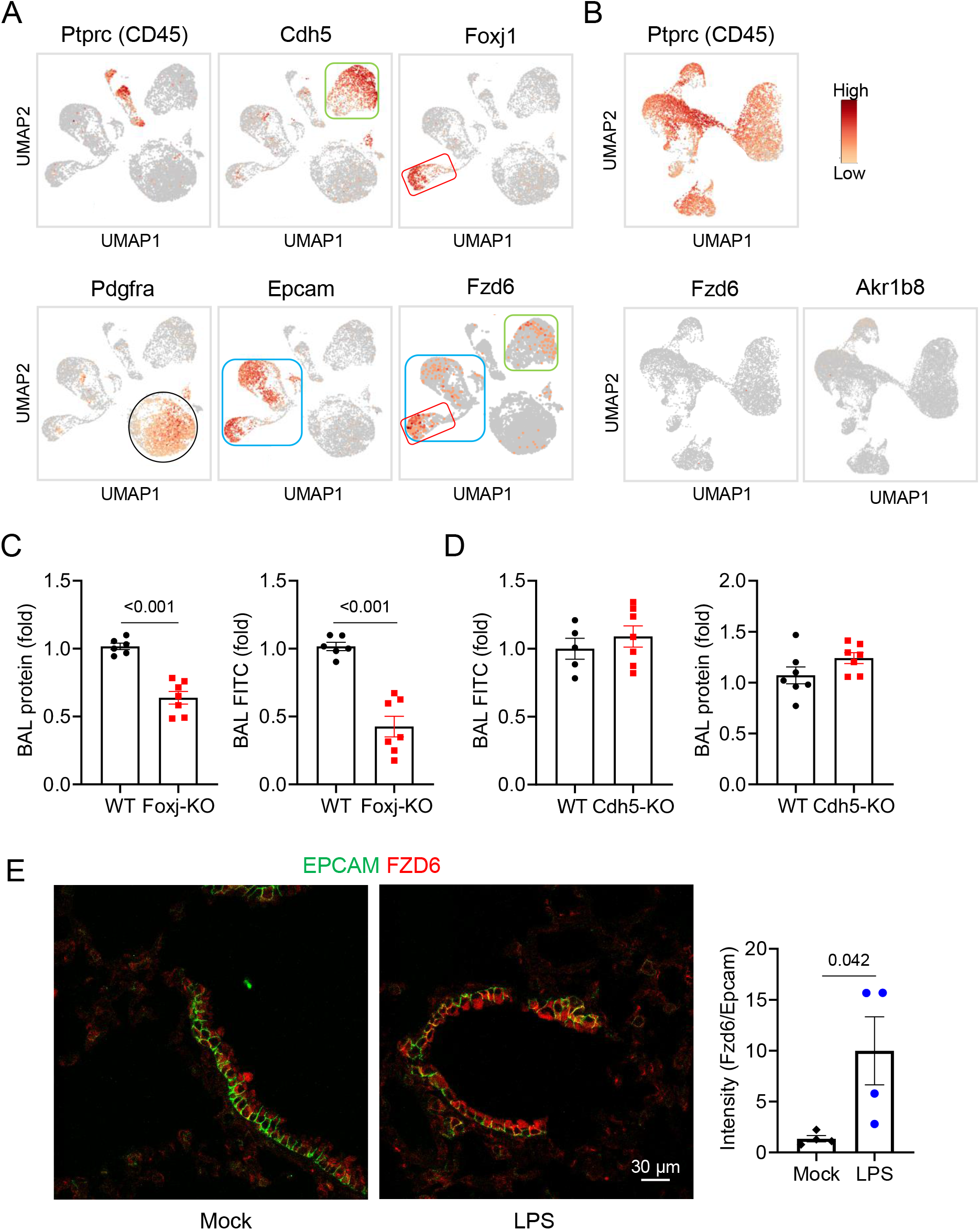
The lack of Fzd6 in lung ciliated epithelial cells enhances pulmonary barrier function. **A-B**. Uniform manifold approximation and projection (UMAP) clustering of lung CD45-negative (**A**) and positive (**B**) cells based on single-cell RNA sequencing data. These cells were sorted after being stained with anti-CD45. Expression of cell type marker genes and Fzd6 are shown. **C-D**. Foxj1-Cre Fzd6 KO (Foxj-KO), Chd5-Cre Fzd6 KO (Cdh5-KO), and their wildtype (WT) controls were treated with tamoxifen and one month later administrated with LPS intratracheally. **E**. Lung sections from mice given vehicle or LPS intratracheally for 18 hours were stained with anti-Epcam and Fzd6, followed by secondary antibodies. The sections were examined under a confocal microscope. The images were quantified by normalizing the area of Fzd6/Epcam double-positive signals to the area of epithelial cells positive signals. Each datum represents one mouse. Five fields were evaluated for each section, and three sections were examined per mouse. Representative sections are shown. Data are presented as means±sem (two-tailed Student’s t-test).

Fzd6 was previously detected by immunostaining at the multiciliated airway epithelial cells [23], which express Foxj1 [24]. Concordantly, we confirmed this observation by detecting colocalization of Fzd6 in the same cells with tdTomato whose expression was driven by Foxj1-Cre (Fig. S2B). Analysis of the expression of all of the ten *Fzd* genes in the Foxj1^+^ cells using our single cell RNA sequencing dataset indicated that Fzd6 expression was the highest (Fig. S2C). Moreover, we found that Fzd6 protein was upregulated in the lungs subjected to LPS injury compared to those without injury (Fig. 2E).

### Fzd6 KO reduces lipid peroxidation in LPS-treated lungs

Consistent with the observations of Mx-Cre-driven Fzd6 KO, there was also an increase in the alveolar number, but not septal thickness, in Foxj1-Cre-driven Fzd6 KO mice over the WT control mice after LPS treatment (Fig. 3A & S3A). The change in alveolar number prompted us to speculate that Fzd6 KO may protect lung cells from alveolar cell death, but we found no significant difference in the number of activated caspase 3-positive cells in the LPS-injured lungs between Fzd6 KO and WT (Fig. S3B). Instead, we observed a decrease in 4-hydroxynonenal (4-HNE, a lipid peroxidation marker) staining in LPS-treated lungs of Fzd6 KO mice compared to those of the WT mice (Fig. 3B-C). In addition, we also detected a decrease in the malonaldehyde (MDA, a byproduct and marker of lipid peroxidation) contents in the BALs of Fzd6 KO mice treated with LPS compared to those of the WT mice (Fig. 3D). Because lipid peroxidation is related to ferroptosis, these data hence suggest that Fzd6 KO may regulate ferroptosis in the LPS-injured lungs.

**Figure 3.**
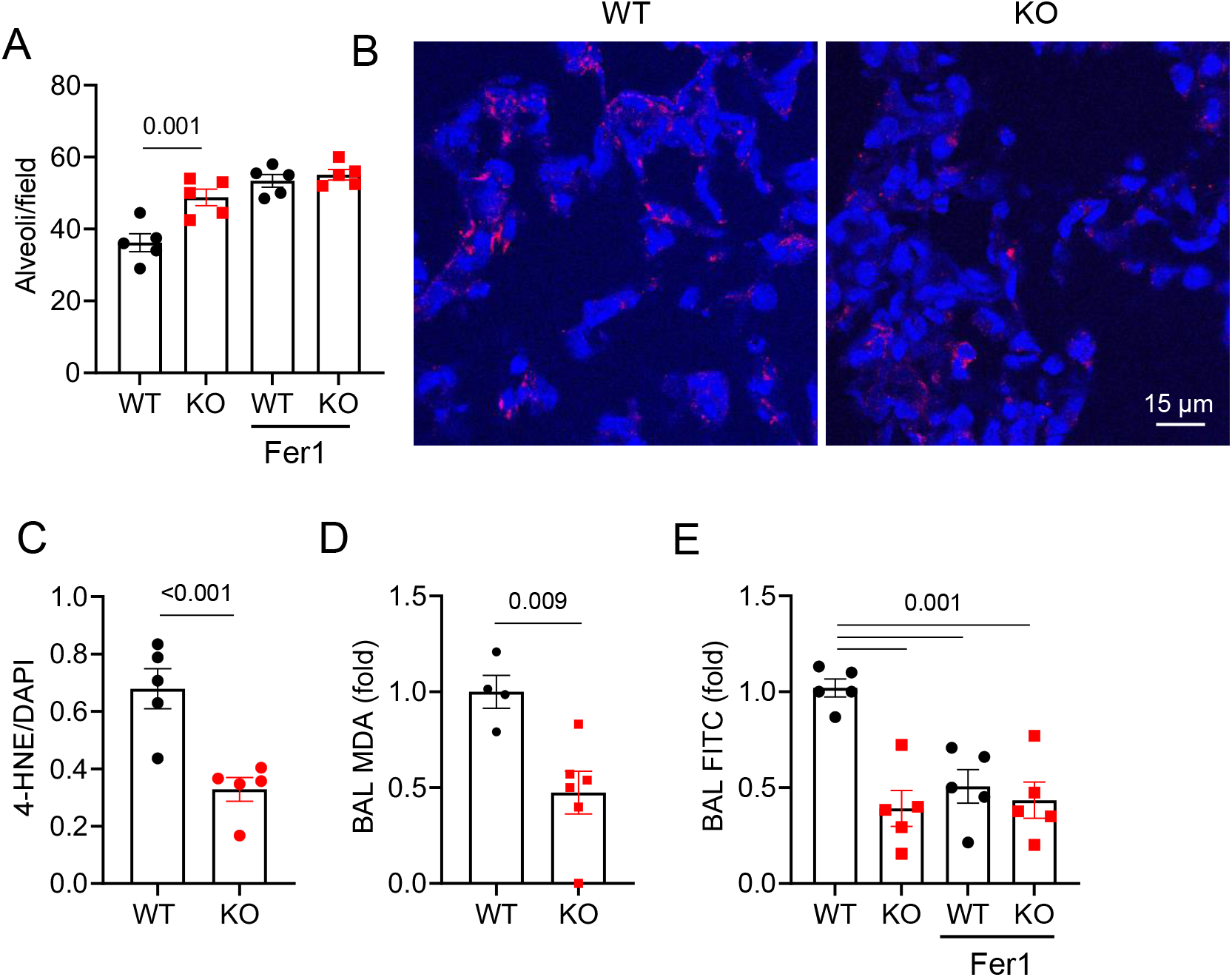
Ferroptosis underlies pulmonary phenotypes of Fzd6 KO. **A,E**. Foxj1-Cre Fzd6 KO and WT mice were treated with LPS as in Fig. 2 and 30 min later administrated via I.V. with mock or ferrostain-1 (Fer1). Lung histomorphometry quantification and permeability measurement were done as in Fig. 1. **B-C**. The lung sections of Foxj1-Cre Fzd6 KO and WT mice treated with LPS were stained with anti-4-HNE (red) and DAPI (blue), and 5 independent sections of each mouse were quantified. Ratios of 4-HNE staining intensity to DAPI intensity are shown in C. Each datum point represents one mouse. **D**. BAL MDA were determined from Foxj1-Cre Fzd6 KO and WT mice treated with LPS. Data are presented as means±sem (two-way Anova for **A,E**; two-tailed Student’s t-test for **C,D**).

To test if ferroptosis may underlie the lung phenotypes of Fzd6 KO, we treated the mice with the ferroptosis inhibitor, ferrostatin-1 (Fer1), which effectively reduced the MDA content in the BALs of LPS-treated lungs (Fig. S3C). Fer1 increased the number of alveoli in LPS-treated WT lungs, but not Fzd6 KO lungs (Fig. 3A). Additionally, Fer1 reduced lung permeability only in WT mice, but not in Fzd6 KO mice (Fig. 3E). Fer1 treatment, like Fzd6 KO, did not affect septal thickness in LPS-injured lungs (Fig. S3A). Because Fer1 did not further alter Fzd6 KO lung phenotypes, it is reasonable to believe that ferroptosis may underlie the lung phenotypes of Fzd6 KO.

### Fzd6 KO suppresses LPS-induced cell death

To directly visualize cell death, we administered povidone iodine (PI) right before mice were euthanized for lung tissue collection. However, we only observed a few PI-positive cells per microscopic view areas at 18 hours post LPS treatment (Fig S4A), which was the end point for the experiments described above. We thus performed a time course and observed more PI-positive cells at 9 hours post LPS treatment than at 18 hours or 4 hours (Fig S4A), suggesting that most of LPS-induced death occurs at a narrow time window. We further examined the PI-positive cells at 9-hour time point and found some of the PI signals were surrounded by CD31 (an endothelial cell marker) staining, whereas others by CD326 (an epithelial cell marker) staining (Fig. 4A). Many of the PI signals were also surrounded by or in vicinity of 4-HNE staining (Fig. 4B). Apoptotic cells were also detected by activated caspase 3 staining at this time. There were about twice number of PI-positive cells than activated caspase3-positive cells (Fig. S4C). Of note, most activated caspase 3-positive cells were PI-negative, suggesting most of apoptotic cells have intact plasma membrane at this time point. Because Fzd6 KO markedly lowered the number of PI-positive cells without affecting the number of activated caspase 3-positive apoptotic cells (Fig. 4C & S4C) and reduced lipid peroxidation (Fig. 3C,D), Fzd6 KO might suppress LPS-induced alveolar cell death primarily via inhibiting ferroptotic cell death.

**Figure 4.**
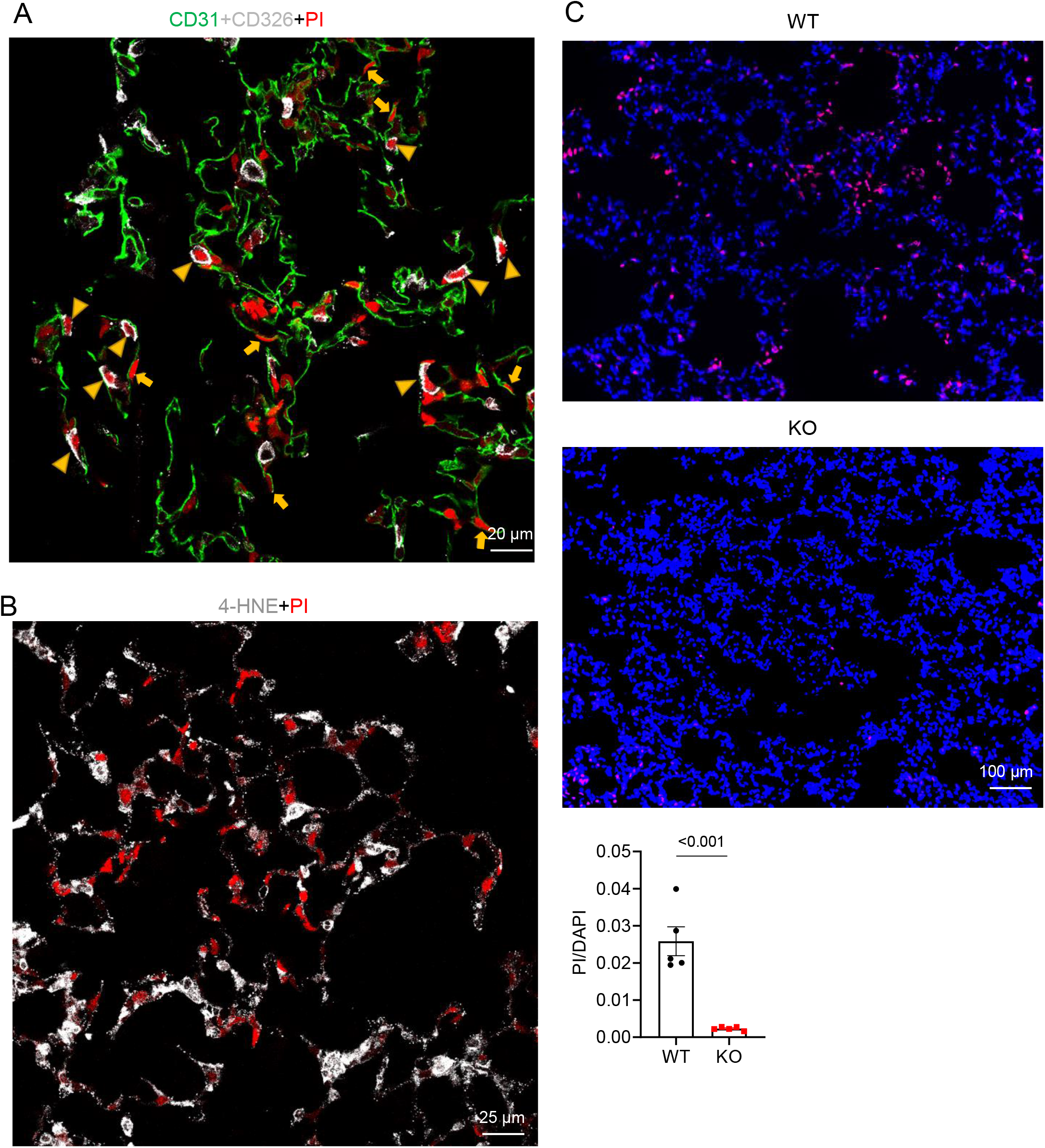
Fzd6 KO suppresses cell death in LPS-treated lungs. **A-B**. A WT mouse was administrated intratracheally with LPS for 9 hours and infused with PI transcardially for 10 min. The lung section was stained with antibodies to CD31, CD326 and 4-HNE. Arrow heads in **A** denote cells double positive for Epcam and PI (dead epithelial cells), whereas arrows denote those double positive for CD31 and PI (dead endothelial cells). **C**. Foxj1-Cre Fzd6 KO and WT mice were treated with LPS as in **A**. The ratios of PI and DAPI signals were quantified from 20 independent sections per mouse lung. Each datum point represents one mouse. Representative fluorescent microscopic images are shown. Data are presented as means±sem (two-tailed Student’s t-test).

### Fzd6 KO increases Ark1b8 expression

To understand how Fzd6 KO regulates LPS-induced lung phenotypes and ferroptosis, we performed single-cell RNA sequencing of CD45-negative cells sorted from Fzd6 KO and WT lungs, respectively. Total 9821 WT cells and 9603 Fzd KO cells passed quality control and were used in the subsequent analysis. Unsupervised clustering yielded 5 major clusters (Fig. S5A). Based on the marker genes expression, epithelial cells are in Clusters 1-3, whereas endothelial cells (EC) are in Cluster 4, and fibroblast cells are in Cluster 5 (Fig. S5B). Additionally, Foxj1-positive ciliated cells are in Cluster 1, whereas Type 2 and 1 alveolar cells are in Clusters 2 and 3, respectively (Fig. S5B). The cell number distributions of these clusters within each genotype are shown in Fig. S5C. Differential gene expression analysis, followed by Ingenuity Pathway Analysis (IPA), revealed enrichment of many pathways in these clusters. The NRF2-mediated oxidative stress pathway and xenobiotic response pathways were the common ones altered in all clusters (Fig. S5D-G). The NRF2 pathway is known to regulate ferroptosis [11-13]. In addition, the ferroptosis pathway was downregulated in the C2+C3 epithelial cell clusters of Fzd6 KO epithelial cells (Fig. S5E) and altered in the C4 endothelial cell cluster (the direction of alteration could be not resolved). These data are consistent with the results that Fzd6 KO affects lipid peroxidation and ferroptosis.

Because Fzd6 KO in the ciliated Foxj1^+^ cells, which line the walls of trachea, bronchioles, and brachiola, reduced alveolar cell death, we suspected this effect of Fzd6 KO was unlikely autonomous. Upon analysis of the differentially expressed genes (DEGs) between Fzd6 KO and WT in the Foxj1^+^ cells, we noticed that Ark1b8 was most significantly upregulated by Fzd6 KO (Fig. 5A & S5H, Table S1). Akr1b8 encodes the aldo-keto reductase family 1 member B8 protein, which reduces aldehydes and their glutathione conjugates. Such enzymatic activities have been implicated to reduce lipid peroxides and hence inhibit ferroptosis [25, 26]. Moreover, Akr1b8 and its human homolog AKR1B10 were found to be secreted from cells through the lysosome-mediated non-classical secretory pathway [27, 28]. The hypothesis that Akr1b8 is a paracrine factor released from ciliated cells is consistent with the single-cell RNA sequencing result that the other common pathways enriched in non-ciliated cells are the xenobiotic metabolism pathways possibly due to Akr1b8’s broad reductase substrate specificity. In supporting the hypothesis, we detected an increased amount of the Akr1b8 protein in the BALs from the Fzd6 KO lungs compared to that from the WT lungs treated with LPS (Fig. 5B). Of note, among all of the highly significantly upregulated or downregulated genes in the Foxj1^+^ cluster, no other genes encode known secreted proteins (Table S1).

**Figure 5.**
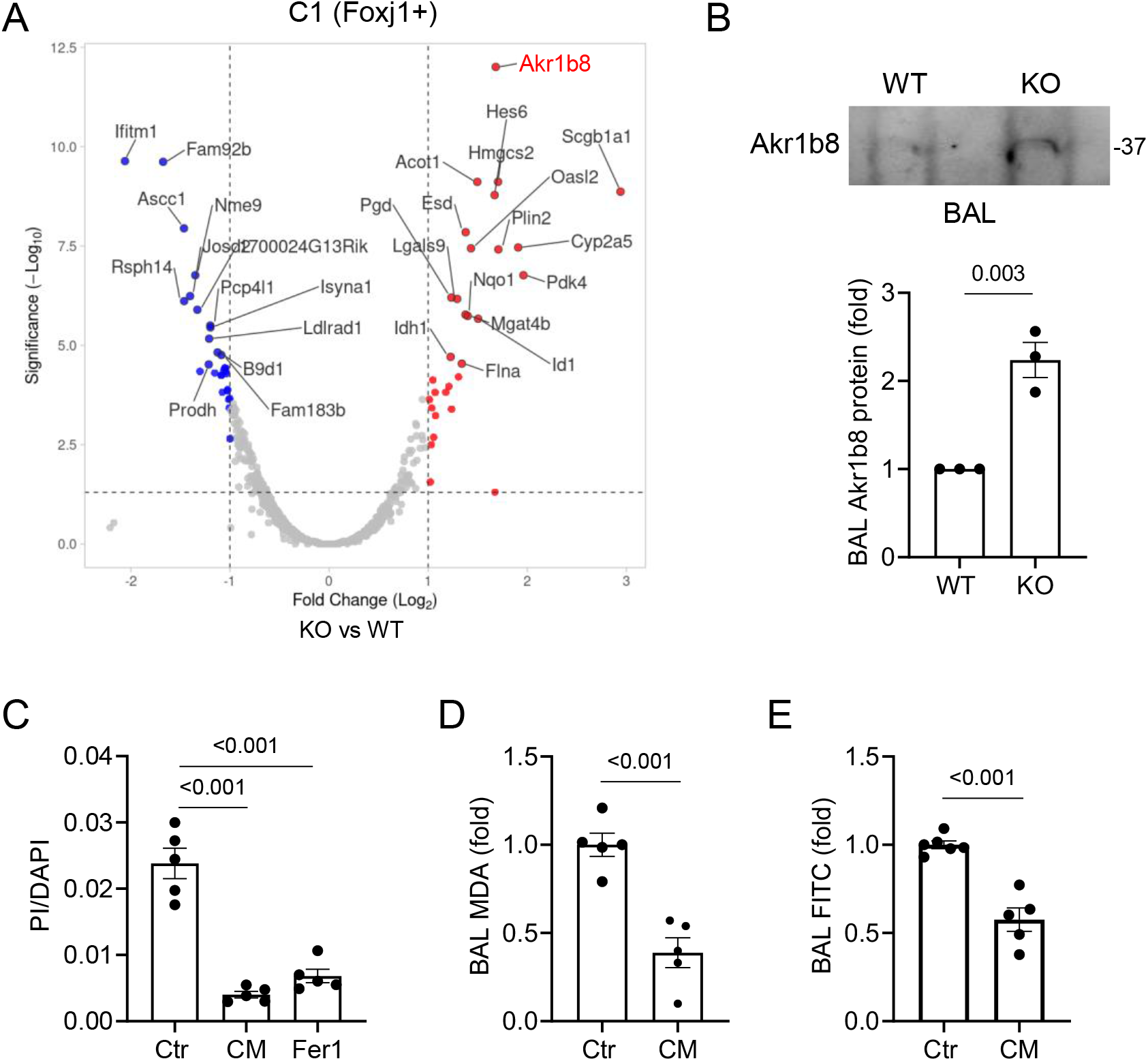
Fzd6 KO increases expression of Akr1b8 to suppress cell death in LPS-treated lungs. **A**. Volcano plot of differentially expressed genes in Foxj1^+^ cells based on single-cell RNA sequencing of CD45-negative cells sorted from Fzd6 KO and WT lungs treated with LPS. **B**. Western analysis of BAL from Foxj1-Cre Fzd6 KO and WT mice given intratracheally with LPS. **C-E**. WT mice were administrated with LPS and PI as in Fig. 4A. The mice were also administrated with either control CM or Akr1b8 CM with LPS. Some of the mice receiving control CM were also given Fer1 intravenously. Samples were collected 9 hours after LPS treatment. The ratios of PI and DAPI signals were quantified and are showed in **C**. BAL MDA and BAL FITC-BSA are shown in **D** and **E**. Data are presented as means±sem (two-tailed Student’s t-test for B, D-E; one-way Anova for C).

### Akr1b8 regulates lung cell death and injury

To test if the Akr1b8 protein has any effect on LPS-induced lung injury, we produced conditioned medium (CM) from HEK293 cells expressing Akr1b8. The Akr1b8 protein was readily detected by Western in the CM (Fig. S5I). We administrated the Akr1b8 CM or control CM via intratracheal instillation 30 minutes after LPS treatment. The Akr1b8 CM markedly reduced the number of PI-positive cells in the lungs (Fig. 5C), the MDA contents in the BAL (Fig. 5D), and pulmonary permeability (Fig. 5E), without significantly affecting BAL neutrophil infiltrates (Fig. S5J). It is worth noting that the effect of the Akr1b8 CM showed the similar efficacy to that of the ferroptosis inhibitor Fer1 in blocking cell death (Fig. 5C).

### Coronavirus lung infection induces ferroptotic alveolar cell death

We next examined if Fzd6 KO could ameliorate lung injury caused by mouse hepatitis virus (MHV), a mouse coronavirus. Foxj1-Cre-driven Fzd6 KO mice showed reduced pulmonary permeability (Fig. 6A) and increased alveolar number (Fig. 6B & S6A) compared to the control mice after intranasal MHV-A59 infection, whereas there were no significant differences in septal thickness (Fig. S6B). There were also markedly reduced PI-positive cells in the lungs of Fzd6 KO mice, accompanied with reduced BAL MDA content (Fig. 6C-D, S6C). In addition, ferroptosis inhibitors Fer1 and lipoxstatin-1 inhibited pulmonary permeability, increased alveolar number, and reduced PI-positive cells in the lungs infected by MHV (Fig. 6E-G) without affecting septal thickness (Fig. S6D). Moreover, neither Fzd6 KO nor ferroptosis inhibitors affected neutrophil infiltration in the MHV-infected lungs (Fig. S6E-F). These results together indicate that Foxj1-specific Fzd6 KO or ferroptosis inhibitors inhibit pulmonary MHV-infection-induced ferroptotic alveolar cell death, which contributes to weakened pulmonary barrier integrity.

**Figure 6.**
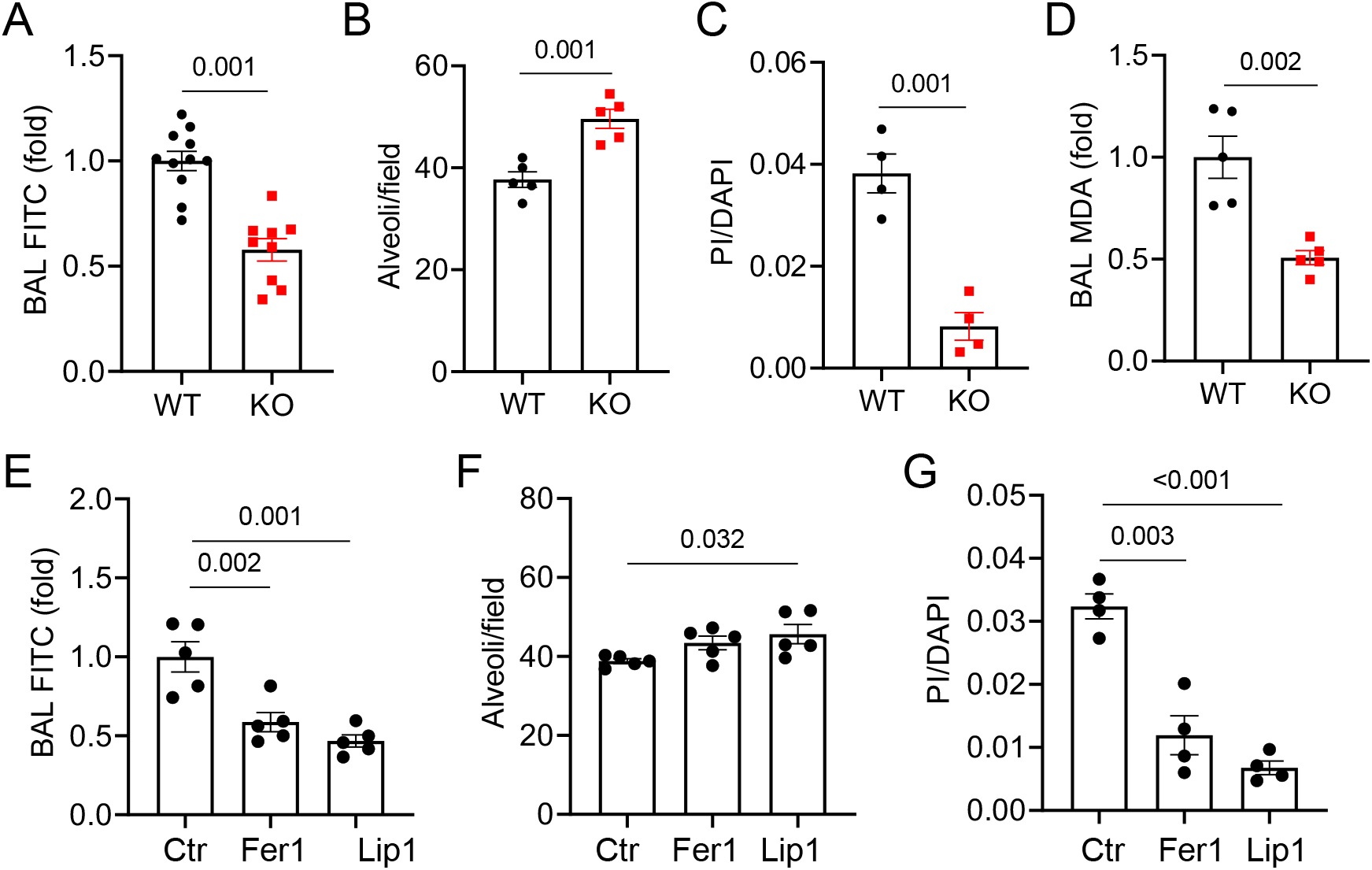
Fzd6 KO and ferroptosis inhibitors enhance barrier function and reduce ferroptotic cell death in MHV-infected lungs. **A-D**. Foxj1-Cre Fzd6 KO and WT mice were infected intranasally with MHV-A59. Two days later, the mice were given FITC-BSA intravenously one hour and PI intracardially 10 min before tissue collection. **E-G**. WT mice were infected as in **A**. They were given one dose of Fer1 intravenously or Liproxstatin (Lip1) intraperitoneally 24 hours later. One day later, the mice were given FITC-BSA intravenously two hours and PI intracardially 10 min before tissue collection. Data are presented as means±sem (two-tailed Student’s t-test for A-D; one-way Anova for E-G).

## DISCUSSION

In this study, we reported that Wnt receptor Fzd6-deficiency in Foxj1^+^ ciliated cells suppressed pulmonary permeability, lipid peroxidation, and alveolar cell death, accompanied with an increase in alveolar number during lung inflammation induced by LPS or coronavirus infection. We also presented evidence to indicate that Fzd6-deficiency suppresses ferroptosis induced by LPS or coronavirus insult via upregulation and secretion of the aldo-keto reductase Akr1b8a from the Foxj1^+^ ciliated cells in the lung. Additionally, we demonstrated that ferroptosis was a major form of death of alveolar cells under these inflammatory conditions that contribute to disruption of pulmonary barrier integrity.

Increases in lipid peroxidation have been shown to occur in lung inflammation and injuries induced by various insult including COVID-19 in animal models [11, 15-17]. We in this report showed for the first time that LPS actually induced a high number of ferroptotic cell death in a narrow window peaked at around 9 hours after the treatment. Because PI-positive cells, which were largely sensitive to Fer1 treatment, were twice more than activated caspase-3-positive cells at this time point, ferroptosis seems to constitute a major and even dominate form of LPS-induced alveolar cell death in WT mice. Interestingly, lipid peroxidation markers were still readily detectable 16-20 hours after LPS treatment. However, very few PI-positive dead cells were detected at this time point. Thus, lipid oxidation markers may not necessarily correlate to ferroptotic cell death.

In this study we also showed that Fzd6 KO in Foxj1^+^ cells, which line the wall of trachea, bronchi, and bronchioles, affects pulmonary permeability and alveolar epithelial and endothelial cell death. In addition, the pathway analysis of DEGs revealed by the single-cell RNA sequencing of WT and Fzd6 KO lung cells showed significant enrichments in a large number of pathways not only in the ciliated cells, but also in other lung cell types including epithelial, endothelial and fibroblast cells. These results suggest the ciliated epithelial cells act on alveolar cells in a non-autonomous mechanism. The single-cell RNA sequencing also uncovered Akr1b8 as the gene upregulated by Fzd6 KO that encodes a putative secreted protein. Akr1b8 and its human close homolog AKR1B10 may be secreted from cells via a noncanonical mechanism [27, 28]. Although the AKR1B molecules are implicated to reduce lipid peroxides and hence inhibit ferroptosis [25, 26], its effect on ferroptotic cell death *in vivo* has not been directly demonstrated. We in this study demonstrated that administration of conditioned medium containing the Akr1b8 protein, but not control CM, could reduce alveolar cell death induced by LPS accompanied by decreasing lipid peroxidation by-product MDA in BALs (Fig. 5). Given Akr1b8 is normally expressed at the highest level in Foxj1^+^ cells (Fig. S5H), the Foxj1^+^ ciliated epithelial cells may act via Akr1b8 to protect alveolar cells from ferroptosis and enhance pulmonary barrier integrity even in the Fzd6 WT lungs. The Fzd6 deficiency in Foxj1^+^ cells merely amplified this interaction between lung ciliated cells and alveolar cells. Therefore, our study revealed a previously unknown role of ciliated cells in protection of alveolar cells from ferroptotic cell death during lung inflammation.

Although Fzd6 expression was detected in other lungs cells, its expression in the Foxj1^+^ ciliated epithelial cells is the highest. Importantly, its expression in the Foxj1^+^ cells is responsible for lung inflammation phenotypes of Fzd6 KO, as Foxj1-Cre Fzd6 KO phenocopied Mx-Cre Fzd6 KO. By contrast, Cdh5-Cre Fzd6 KO did not affect these phenotypes. In addition, Fzd6 expression was upregulated upon LPS treatment. Thus, it seems that Fzd6 constitutes a pro-ferroptotic mechanism during acute lung inflammation. It is rather difficult to envision how this pro-ferroptotic role of Fzd6 would be beneficial for the normal physiology of the organism. We thus speculate that this role of Fzd6 is more likely a by-product of normal biological function of Fzd6. Fzd6 has an important role in establishment of planar cell polarity during development as its deficiency disrupts axon growth and guidance, hair follicle orientation, and stereociliary bundle orientation in inner ear sensory hair cells [29-31]. Fzd6 was also implicated in regulating airway cilial orientation, which is important for concerted, directional cilial motility and proper mucociliary clearance [23]. It would be interesting to determine if conditional inactivation of Fzd6 in adult would also cause planar polarity defects in future studies. Another question that remains to be investigated is how Fzd6 signaling represses Akr1b8 expression in ciliated cells. The pathway analysis of the single-cell sequencing data did not implicate Wnt signaling pathways among those altered by Fzd6 KO in ciliated cells. Future studies need to further address this question.

## Materials and Methods

### Mice

Fzd6^fl/fl^ (*Fzd6*^*tm2*.*1Nat*^/J) and Foxj^CreER2^ (*Foxj1*^*tm1*.*1(cre/ERT2/GFP)Htg*^/J) mice were obtained from the Jackson Laboratory. Akr1b8+/-mice (C57BL/6N-*Akr1b8*^*tm1*.*1(KOMP)Vlcg*^/JMmucd) were obtained from MMRRC. MX1^Cre^ and Cdh5^CreER2^ were described previously [32, 33]. All mice used in this study were back-crossed with the C57BL/6J background. Fzd6 gene disruption was induced by intraperitoneal injection of 40 ul of poly-I:C (10 mg/ml) into the MX1^Cre^ mice every other day for four treatments or by intraperitoneal injection of 100 ul tamoxifen (20 mg/ml) for 6 consecutive days into the FoxJ^CreER2^ and Cdh5^CreER2^ mice. All animal experiments were performed with the approval of the Institutional Animal Care and Use Committee at Yale University.

### LPS-induced lung inflammation

Mice were anesthetized with ketamine/xylazine (100 and 10 mg/kg). A 22-gauge catheter (Jelco, Smiths Medical) was guided 1.5 cm below the vocal cords, and LPS (50 μl, 1 mg/ml, Escherichia coli 011:B4) was instilled while mice postures were maintained upright. Sixteen hours after the induction of injury, 100 μl of fluorescein isothiocyanate–labeled albumin (FITC-BSA, 10 mg/ml) was injected into mice via retro-orbital vein. Mice were then euthanized for sample collection 1-2 hours later. BALs were collected after 1 ml of PBS was instilled into lungs and retrieved via a tracheal catheter. For the baseline permeability measurement, saline without LPS was administered the same way. The baseline permeability measurement was subtracted in the data presented.

For ferrostatin-1 (Fer1) treatment, mice were administrated with Fer1 (1.0 mg/kg) via retro-orbital vein one hour after LPS instillation. Fer1 was dissolved in DMSO at the concentration of 200 mg/ml and diluted to a concentration of 0.2 mg/ml in PBS.

### Histology

The right ventricles were perfused with 10 ml of PBS, and the lungs were inflated with 4% paraformaldehyde at a constant pressure of 25 cm H^2^O. The lungs were then fixed in 4% paraformaldehyde for 24 h at 4°C. The lung tissues were embedded in paraffin. They were cut into 5 μm-thick sections and stained with H&E at the Comparative Pathology Research Core at Yale School of Medicine. The sections were imaged using the Keyence BZ-X800 microscope. For quantification of the histological images, 20 fields per mouse were selected via random sampling [34]. For each field, a 42-point lattice with grid line was used to perform quantitative microscopy as described in [35, 36]. Measurements were normalized to mouse body weight. All measurements were done by two blinded observers.

### Immunohistochemistry

Lungs were perfused, inflated, and fixed with 4% PFA for 4-6 h on a shaker at 4°C. They were then washed with PBS three times and perfused in 30% sucrose solution in PBS overnight at 4°C. They were subsequently mounted in OCT embedding compound and frozen first at −20°C and then at −80°C. Tissue sections were prepared at 8 μm thickness with a cryostat and mounted onto gelatin-coated histological slides, which were stored at −80°C. For immunostaining, slides were thawed to room temperature, fixed in pre-cooled acetone for 10 min, and then rehydrated in PBS for 10 min. The slides were incubated in a blocking buffer (1% horse serum and 0.02% Tween 20 in PBS) for 1 h at room temperature, then incubated with antibodies diluted in the blocking buffer overnight at 4°C. The slides were then washed three times with PBS and incubated with a secondary Ab in the incubation buffer for 1 h at room temperature. After repeated washes, the slides were mounted with an anti-fade mounting medium containing DAPI (Thermo Fisher Scientific, P36931) and visualized with confocal microscope (Leica SP5). Fluorescence images were quantified by using ImageJ software. Antibodies used are: anti-Fzd6 (Novus, AF1526), anti-CD326 Abs (Invitrogen, MA5-12436), anti-CD31 (BioLegend 102421), anti-4-HNE (Abcam ab46545), and anti-activated caspase 3 (ThermoFisher 560627).

### Flow cytometry analysis of BAL and lung-infiltrating leukocytes

Leukocytes in BAL were pelleted by centrifugation (500 × g for 5 min at 4°C), followed by staining and flow cytometry. To prepare lung-infiltrating leukocytes, mouse right ventricles were perfused with 10 ml of PBS, excised, gently disrupted by teasing, and resuspended in RPMI-1640 medium supplemented with 250 U/ml of collagenase, 400 U/ml elastase and 50 U/ml of DNases. Lungs were then enzymatically digested for 40 min at 37°C. Dispersed cells were filtered through a 70-μm cell strainer to eliminate clumps and debris. After centrifugation for 5 min (500 × g) at 4°C, cell pellets were resuspended in RBC lysis buffer (Sigma-Aldrich, R7757) and incubated at room temperature for 5 min to remove erythrocytes. Cells were pelleted again and resuspended in PBS, fixed with 2% PFA (Santa Cruz, sc-281692), and stained with antibodies for 1 h on ice in the dark.

Antibodies used for flow cytometry are as follows: mouse CD45-BUV395 (BD Biosciences), mouse Ly6G-Pacific Blue ([BioLegend](http://www.biolegend.com/)), and mouse Ly6C-FITC ([BioLegend](http://www.biolegend.com/)).

### BAL Cytokine measurement (ELISA)

BALs were centrifuged at 800Xg at 4°C for 10 min to collect cell-free BAL supernatants. The levels of IL-6, IL-1b and TNF-a were determined by using enzyme-linked immunosorbent assay (ELISA) kits specific for the cytokines according to the manufacturer’s instructions (Proteintech).

### BAL MDA measurement

BAL MDA contents were determined using the MDA adduct competitive ELISA kit (Cell Biolabs, Cat#: STA-832) according to the manufacturer’s instructions.

### *In vivo* propidium iodide (PI) staining

Mice were perfused transcardially with PI (Sigma-Aldrich, P4170, 0.75 mg/ml, 4ml/mice in PBS) for 10 min. The lungs were then fixed by perfusing transcardially with 4 ml 4% PFA (containing 0.5 mg/ml PI) 10 min. Lung architecture was preserved by inflation the lung with OCT solution (OCT:PBS = 1:1), followed by additional fixation with 4% PFA for overnight. The lungs were then transferred to 30% sucrose and incubated for additional 24 hrs. Finally, the lung samples were embedded in OCT and stored at −80°C until sectioning. Sections (10 μm) were counterstained with DAPI and examined under a fluorescent microscope.

### Akr1b8 conditional medium preparation

The cDNA for mouse AKR1b8 was obtained from Horizon (Cat# MMM1013-202763109) and subcloned to a mammalian expression vector. HEK-293T cells were maintained in DMEM supplemented with 10% FBS at 37oC and transfected with Akr1b8 expression plasmid or empty vector using Lipofectamine LTX. After overnight, the cells were washed three time with PBS and then replaced with 100 mL of fresh DMEM media without FBS. After 24 h, the conditioned media (CMs) were collected and centrifuged at 15000×g for 5 min at room temperature. The CMs were concentrated with Amicon Ultra-15 centrifuge tubes (Merck Millipore Ltd., Saint-Quentin-Fallavier, France) by 10 folds. The CMs were then diluted 30 folds into LPS for intratracheal instillation.

### MHV-A59 lung infection model

Mice were anesthetized with ketamine/xylazine and instilled with 30 ul PBS containing MHV-A59 intranasally. For ferroptosis inhibitor treatment, mice were injected with Fer1 (1.0 mg/kg) via retro-orbital vein or Liproxstatin-1 (8 mg/kg) intraperitoneally.

### Single-cell RNA sequencing and analysis

Mouse right ventricles were perfused with 10 ml of PBS, excised, gently disrupted by teasing and resuspended in RPMI-1640 medium supplemented with 250 U/ml of collagenase, 400 U/ml elastase and 50 U/ml of DNases. Lungs were then enzymatically digested for 40 min at 37°C. Dispersed cells were filtered through a 70-μm cell strainer to eliminate clumps and debris, and erythrocytes were lysed using the lysing buffer. Cells in single-cell suspension were resuspend cells in cold 0.1% BSA/PBS. Following live/dead staining with viability dye (Fixable Viability Dye eFluor 506, eBioscience), cells were incubated with a Fc-blocking reagent (BD Biosciences) for 5 minutes at 4 °C and then incubated with an anti-CD45.2 mAb-PE-Cy7 antibodies for 1 hour at 4°C. Cells were sorted using a 100 μm nozzle and 40 psi pressure (FACSAria instrument, BD Biosciences). Single-cell 3’ RNA-seq libraries were prepared using the Chromium Single-cell V3 Reagent Kit and Controller (10x Genomics). Libraries were assessed for quality control and then sequenced using a HiSeq 4000 instrument (Illumina). The demultiplexing, barcoded processing, gene counting, and aggregation were performed using the Cell Ranger software v3.1.0 (https://support.10xgenomics.com/single-cell-gene-expression/software/pipelines/latest/what-is-cell-ranger) with reference transcriptome GRCh38. Further downstream analysis was performed in Seurat V4.3.0 [37]. Initially, cells were chosen based on unique feature counts less than 2500 or greater than 200 and mitochondrial read counts <5% and data is log normalized. After filtering the cells, UMI counts were variance-stabilized using scTransform with 3000 most variable features. Top 30 principal components were used to construct SNN graph and embedding with UMAP. Major cell types were detected using expression of key cell type markers across clusters. Differential gene expression analysis was performed between two conditions and fold change were measured. All raw and processed data of this study were deposited at Gene Expression Omnibus GSE GSE222810.

## FIGURE LEGENDS

**Figure S1.**
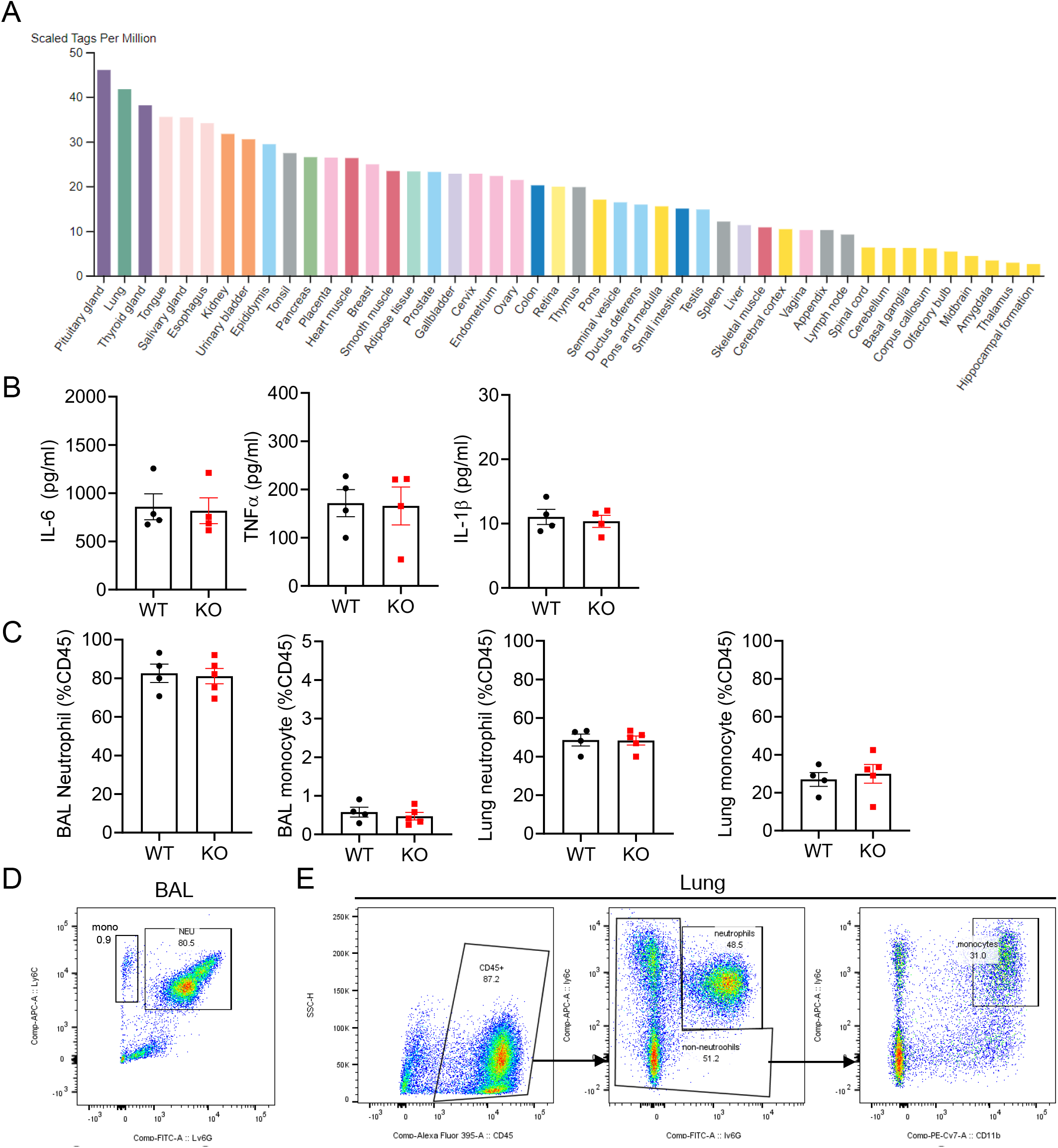
Fzd6 KO alters responses to pulmonary inflammation induced by LPS. **A**. Fzd6 expression data in different human tissues generated by the FANTOM5 project using the Cap Analysis of Gene Expression (CAGE). **B**. BAL cytokines were determined by ELISA. **C**. The abundance of BAL and lung myeloid cells were analyzed by flow cytometry. **D-E**. Flowcytometry gating strategies for BAL (**D**) and lung digest (**E**). Data are presented as means±sem.

**Figure S2.**
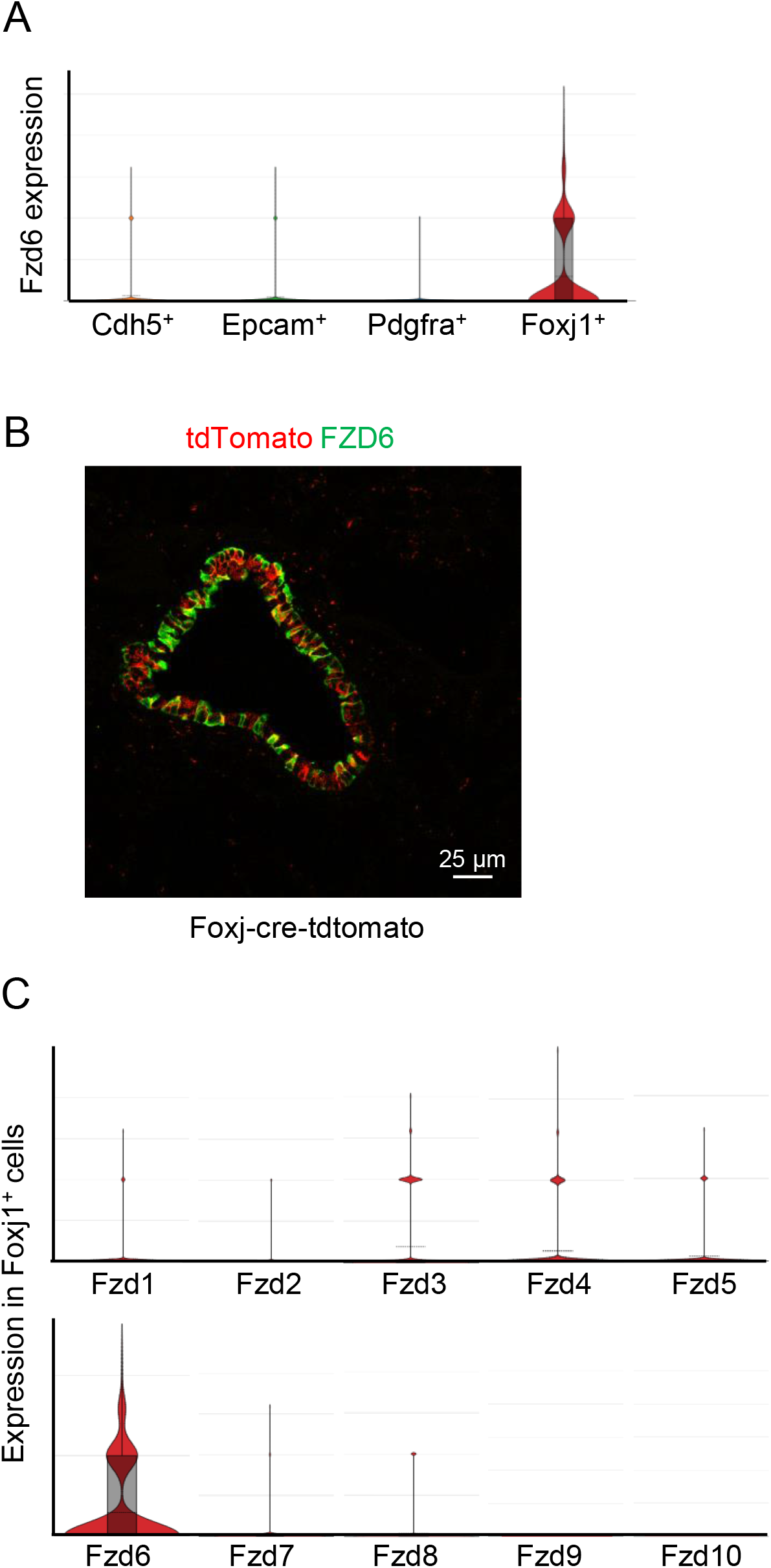
Expression of Fzd6 in lung ciliated epithelial cells. **A**. Expression of Fzd6 in the four clusters shown in Fig. 2A. **B**. A lung section from a Foxj1-Cre Tdtomato mouse was stained with the anti-Fzd6 antibody. **C**. Expression of all of the ten Fzd genes in the Foxj1^+^ cluster.

**Figure S3.**
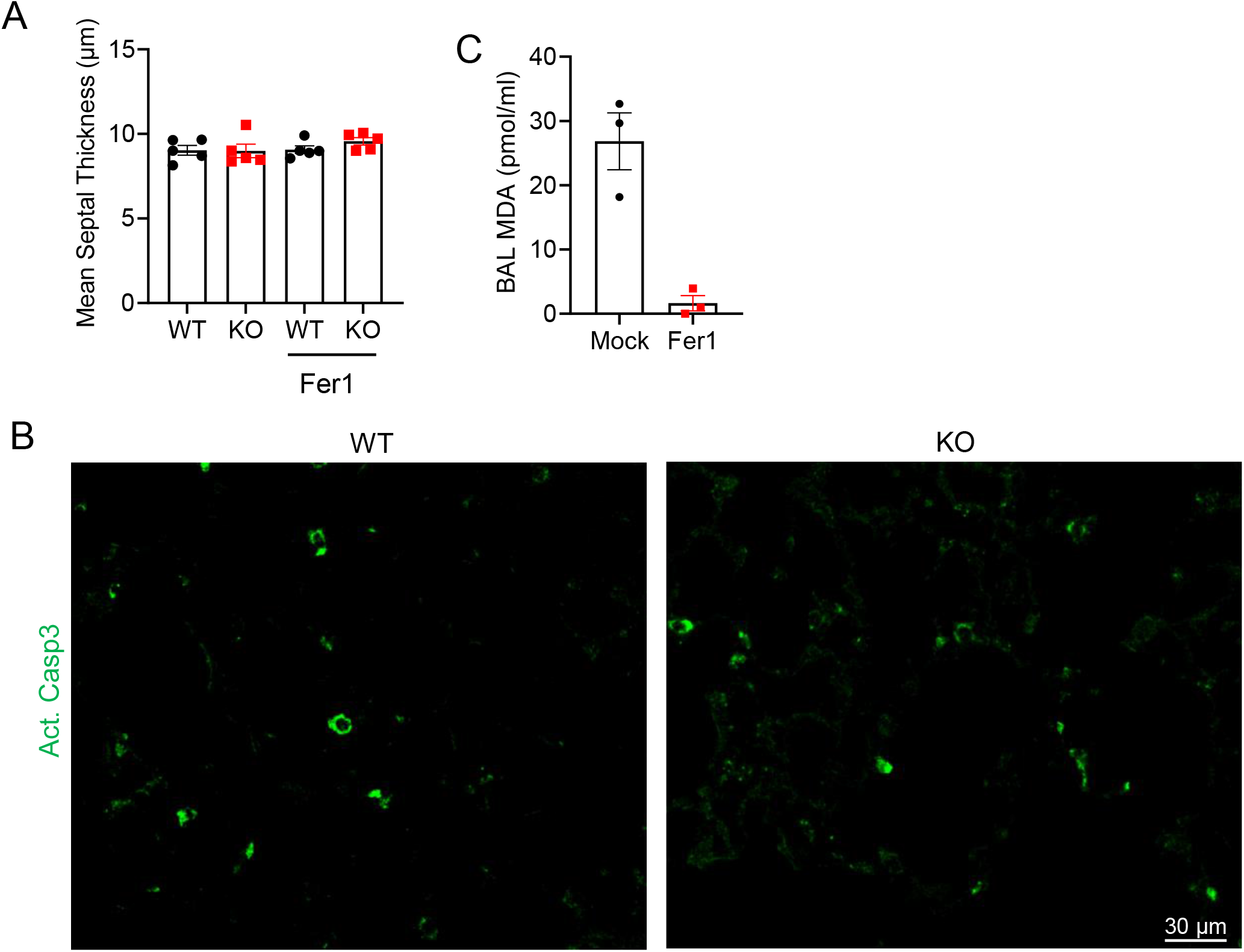
Ferroptosis is important for pulmonary phenotypes of Fzd6 KO. **A**. Additional histological section quantifications for Fig. 3A. **B**. WT and Foxj1-Cre Fzd6 KO mice were administrated intratracheally with LPS. Lung sections were collected after 18 hours and stained with anti-activated caspase3 (Act. Casp3) antibody. **C**. The WT mice were administrated intratracheally with LPS and intravenously with mock or Fer1. BAL MDA was determined 18 hours later. Data are presented as means±sem (two-tailed Student’s t-test).

**Figure S4.**
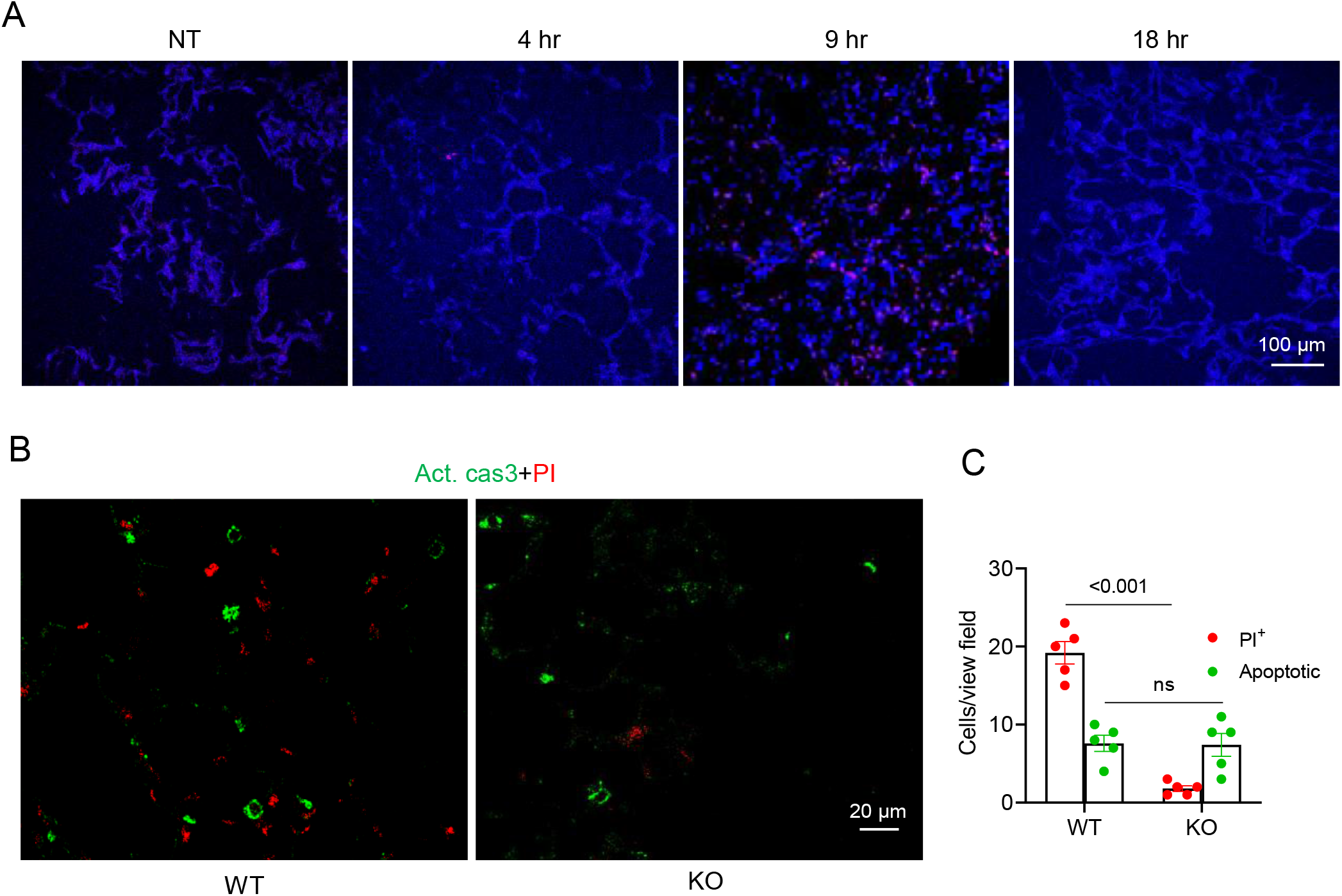
LPS-induced lung cell death peaks at 9 hours after the treatment. **A**. WT mice were administrated intratracheally with LPS for different times and infused with PI 10 min before sample collection. **B-C**. the lung sections described in Fig. 4A were stained with anti-activated caspase 3 (act. Cas3). PI and act. Cas3-positive cells were quantified from 5 independent sections per mouse lung. Each datum point represents one mouse. Data are presented as means±sem (two-tailed Student’s t-test).

**Figure S5.**
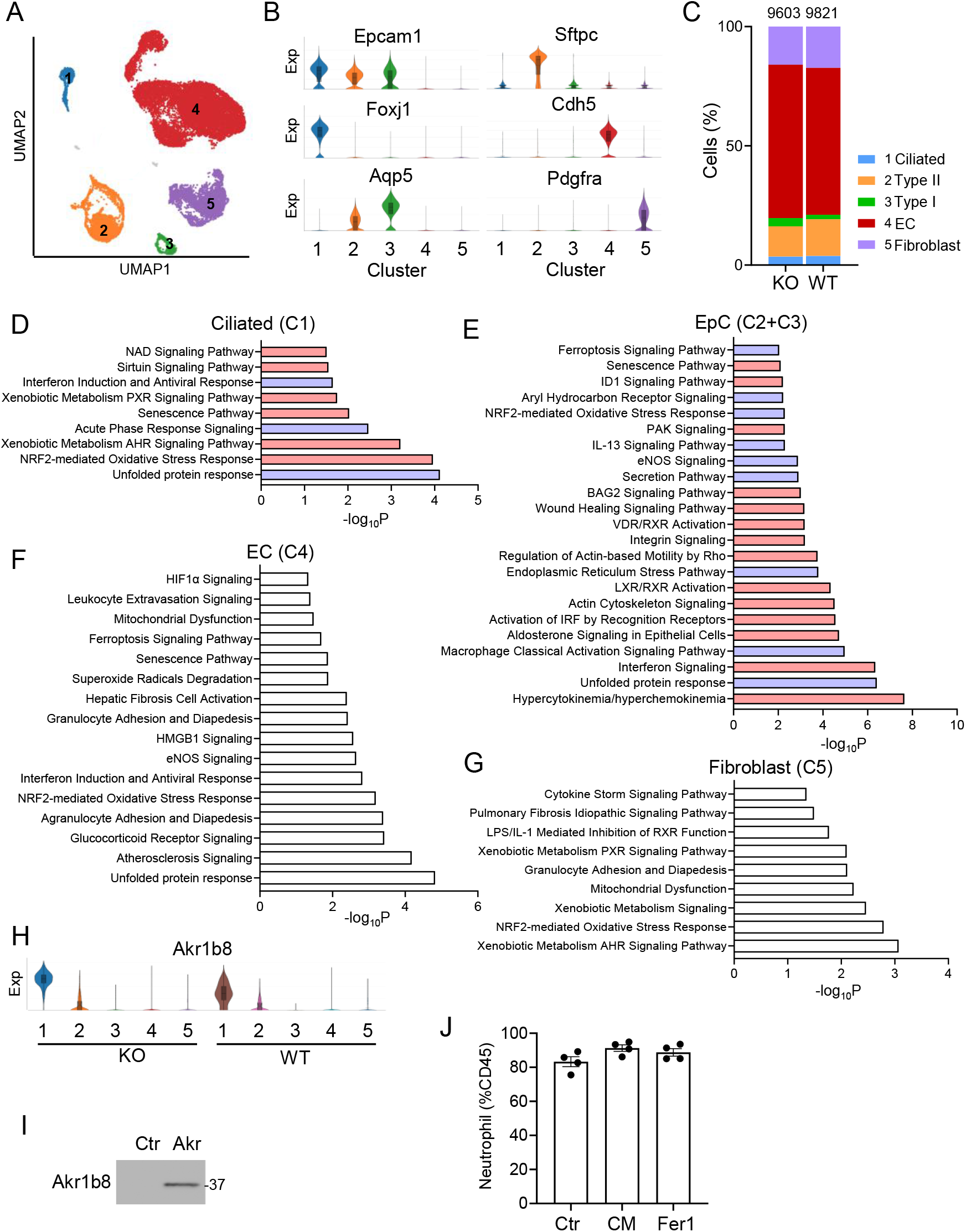
Single cell RNA sequencing of Fzd6 KO and WT lung cells. **A**. UMAP clustering of Lung CD45-negative lung cells sorted from two pairs of WT and Fzd6 KO mice. **B**. Cell type marker gene expression (Exp) of the clusters. **C**. Cell number proportion of each cluster. **D-G**. Ingenuity Pathway Analysis (IPA) of DEGs in the clusters. Pink columns represent upregulated pathways in Fzd6 KO, whereas blue columns represent downregulated ones. Open columns represent undetermined regulation direction. **H**. Akr1b8 expression in the 5 clusters classified in **A** of WT and Fzd6 KO samples. **I**. Western detection of the Akr1b8 protein in conditioned medium used in Fig. 5. **J**. Neutrophil abundance in the BALs of mice treated with control CM (Ctr), Akr1b8 CM (CM) or ferrostatin-1 (Fer1).

**Figure S6.**
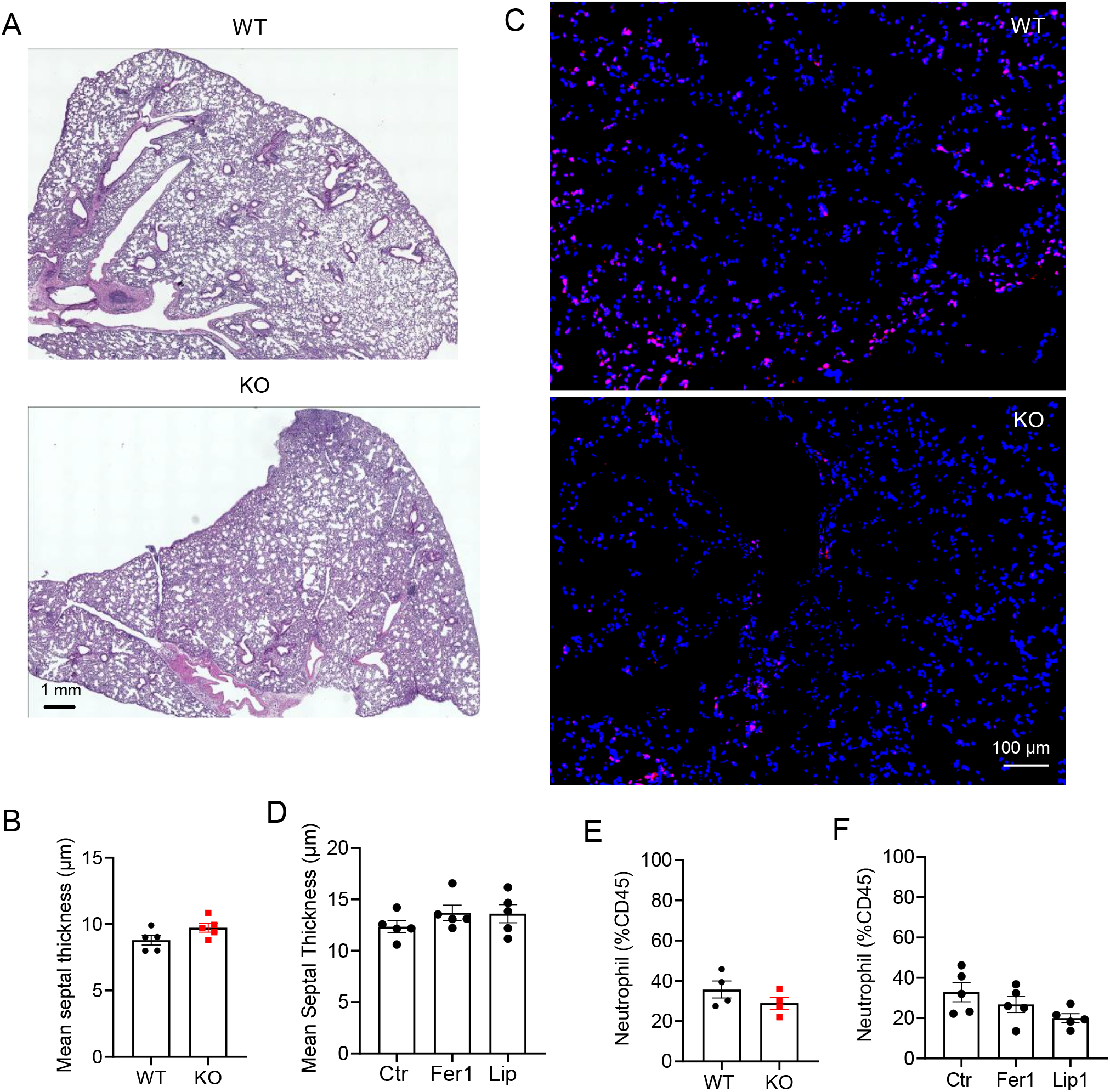
Effects of Fzd6 KO or ferroptosis inhibitors on MHV-59A infected lungs. **A-B**. Representative images for Fig. 6B and additional lung histomorphometry quantification. **C**. Representative images for Fig. 6C. **D**. Additional lung histomorphometry quantification for Fig. 6F. **E**. BAL neutrophils in the lungs described in Fig. 6A were analyzed by flow cytometry. **F**. BAL neutrophils in the lungs described in Fig. 6F were analyzed by flow cytometry. Data are presented as means±sem.

